# Optimal inventorying and monitoring of taxonomic, phylogenetic and functional diversity

**DOI:** 10.1101/060400

**Authors:** Pedro Cardoso, Miquel A. Arnedo, Nuria Macías-Hernández, William D. Carvalho, José C. Carvalho, Renato Hilário

## Abstract

Comparable data is essential to understand biodiversity patterns. While inventorying requires comprehensive sampling, monitoring focuses on as few components as possible to detect changes. Quantifying species, their evolutionary history, and the way they interact claims for studying changes in taxonomic (TD), phylogenetic (PD) and functional diversity (FD). Here we propose a method for the optimization of sampling protocols for inventorying and monitoring diversity across these three diversity dimensions taking sampling costs into account.

We used Iberian spiders, Amazonian bats and Atlantic Forest mammals as three case-studies. The optimal combination of methods for inventorying and monitoring required optimizing the accumulation curve of α-diversity and minimizing the difference between sampled and estimated β-diversity (bias), respectively.

For Iberian spiders, the optimal combination for TD, PD and FD allowed sampling at least 50% of estimated diversity with 24 person-hours of fieldwork. The optimal combination of six person-hours allowed reaching a bias below 8% for all dimensions. For Amazonian bats, surveying all the 12 sites with mist-nets and 0 or 1 acoustic recorders was the optimal combination for almost all diversity types, resulting in >89% of the diversity and <10% bias with roughly a third of the cost. Only for phylogenetic α-diversity, the best solution was less clear and involved surveying both with mist nets and acoustic recorders. For Atlantic Forest mammals the optimal combination to assess all types of α- and β-diversity was to walk all the 10 transects and no camera traps, which returned >95% of the diversity and <5% bias with a third of the costs.

The widespread use of optimized and standardized sampling protocols and regular repetition in time will radically improve global inventory and monitoring of biodiversity. We strongly advocate for the global adoption of sampling protocols for both inventory and monitoring of taxonomic, phylogenetic and functional diversity.

## Introduction

The current level of biodiversity loss is, so far, incommensurable, with our era being recognized as the grand stage to the sixth extinction era (Leakey & Lewin, 1996; Cowie et al., 2022). Available evidence suggests that probably more species go extinct daily than they are described. Yet, useful knowledge for conserving most species is extremely scarce and the current population trends for most taxa are unknown (Cardoso et al., 2011a, 2020). Currently, information about biodiversity is highly concentrated both geographically and taxonomically (Cardoso et al., 2011a; Hortal et al., 2015; Mammola et al., 2023). Improving our knowledge of global biodiversity and assessing its changes is a huge challenge, especially considering that most of the biodiversity occurs in developing countries, where there are fewer resources (both human and financial) to carry out inventories and monitoring (Rodrigues et al., 2010; Hortal et al., 2015). Therefore, there is a need to optimize sampling protocols in order to better allocate sampling effort and resources (Gardner et al., 2008; Cardoso & Leather, 2019).

### Inventorying or monitoring?

The answer to this question is most of the time obvious. Although many researchers and practitioners tend to see these tasks as equivalent or interchangeable, the processes and goals are essentially different. An inventory is required if the goal is to know which species, clades or traits live where, to study community assemblage patterns and its drivers or to prioritize areas for conservation, among many other uses that require comprehensive knowledge of community composition. As the aim is to maximize sampled α-diversity, all taxa of interest should be targeted, a high level of sampling completeness is needed, and sampling should cover as much as possible the entire region and seasons (Table 1). On the other hand, this is either a one-time or a rare event that given the effort involved is almost invariably impossible to replicate in the short or even medium-term for all but species-poor taxa.

**Table 1.** Characteristics of inventorying and monitoring of biodiversity.

|  | inventorying | monitoring |
| --- | --- | --- |
| **Target** | maximize sampled α-diversity | minimize difference between sampled and true β-diversity |
| **Taxonomic scope** | All taxa | Selected taxa |
| **Sampling effort** | Major ($\geq 50\%$ completeness) | Minor (often $< 50\%$ completeness) |
| **Geographic scope** | Global | Selected sites |
| **Seasonality** | Full year | Selected season (often of maximum diversity) |
| **Periodicity** | Single or sporadic ($> 10$ years) | Regular (1 to 10 years) |

In contrast, monitoring is required if the goal is to detect, and if possible quantify, changes in community composition (or of a single species of particular interest) in space or, more often, over time (Noss, 1990; Stork et al., 1996; Bawa & Menon, 1997; Hilty & Merenlender, 2000; Yoccoz et al., 2001; Danielsen et al., 2005). To successfully detect changes in community composition, i.e. β-diversity, one should, at any sampling occasion, minimize differences between sampled and true (or estimated) β-diversity. To achieve this, one may focus on selected taxa, which means that completeness is not required, and sampling can cover only a subset of sites sampled in the height of seasonal abundance for the selected taxa (Table 1). This minimizes the required time and resources and allows frequent replication. In other words, if inventorying often requires comprehensive, time-consuming methods, monitoring should build on it and focus on as few components as possible that still allow the correct perception of changes on predetermined indicators in time, or often, space (Table 1; Stork et al., 1996; Longino & Colwell, 1997). To sample or acquire useful information for both purposes can therefore be viewed as fundamentally different but complementary processes, herewith called α and β-sampling (Table 1).

Both inventorying and monitoring of biodiversity require defining what biodiversity is in the first place. Its usual meaning covers the variety of species, their evolutionary history and how they interact to constitute complex assemblages of communities and ecosystems. Quantifying all these aspects leads to the need to study three different facets of biodiversity: taxonomic (TD), phylogenetic (PD), and functional (FD) diversity, also covering species, genes and ecosystems (DeVictor et al., 2010). The existence of optimized protocols to inventory and monitor TD, PD and FD of a much wider range of taxa is therefore essential if we really want to follow the current trends in biodiversity as a whole, from local to global scales (Cardoso & Leather, 2019). Several global initiatives are currently underway for the inventory and monitoring of biodiversity (Jetz et al., 2019). Monitoring has however been the main concern, given the current rates of biodiversity loss. Some of the heavy users of monitoring data are the GEO BON - Group of Earth Observation Biodiversity Observation Network, within which Pereira et al. (2010) proposed a global terrestrial species monitoring program, advocating that vascular plants and birds should be the candidate taxa for that purpose. Pereira et al. (2013) noted that Essential Biodiversity Variables relating to species, genes or traits require representative sampling across taxonomic groups, recognizing the difficulty in obtaining such data. The GEO BON set of variables to follow is now expanding to include other taxa such as butterflies (van Swaay et al., 2015). Furthermore, the IUCN (International Union for the Conservation of Nature) proposed the Red List Index (RLI), which applies to any organism group, yet only calculated for a handful of taxa (Butchart et al., 2010; Henriques et al. 2020). Finally, the Living Planet Index (LPI) is based only on vertebrate population trends, limiting its application and interpretation to most of the global biodiversity, i.e., invertebrates and plants (Ledger et al., 2023). All these global initiatives have several points in common. They do not currently consider most taxa, and there is no evidence that the ones covered can serve as good surrogates for detecting trends in the remaining biodiversity.

The lack of comparable data is in part due to the very nature of species-rich taxa. The sheer diversity of invertebrates requires novel approaches to collecting data useful to track current changes (Van Klink et al., 2022). But even large and species-poor taxa might be difficult to track in regions with both high diversity and little resources. Cardoso and Leather (2019) proposed a general framework to tackle this problem by eliminating biases and optimizing biodiversity inventorying and monitoring. Also, Cardoso (2009) proposed a method to optimize the inventory of species (TD) (COBRA protocol: Conservation Oriented Biodiversity Rapid Assessment), particularly focusing on mega-diverse groups and using spiders as a case study. This former study demonstrated that it is possible to sample in a standardized, yet optimized way, and the protocol proposed just a decade ago is now being used worldwide (Emerson et al., 2017; Malumbres-Olarte et al., 2017; Crespo et al., 2018; Kiljunen et al., 2020). Here we extend the concept from inventorying to monitoring and its application from TD to PD and FD. We also propose how to take into account different costs, both fixed and variable, implied by different sampling methods. We demonstrate how to optimize sampling protocols for both invertebrates (Iberian spiders) and vertebrates (Amazonian bats and Atlantic Forest medium and large-sized mammals).

## Materials and methods

### Basic algorithm for inventorying (α-sampling)

In an ideal world, sampling biodiversity would imply no costs. In the real world, any sampling event has two kinds of costs. Fixed costs imply some kind of initial investment, such as buying traps or paying trips. Variable costs add up with the sampling effort, for example paying person-hours of field or lab work or buying extra tubes and reagents to store and process samples. All these should be taken into account for any cost-benefit analysis. In addition, we can build on a predetermined set of samples to improve their efficiency with complementary methods. This constrained algorithm may be useful when we want to improve on an existing sampling protocol, guaranteeing a minimum common denominator to allow comparisons between old and new protocols. Improving a protocol based on preexisting samples implies forcing the optimization exercise to start with such constraints and searching for the methods that better complement it when adding further samples (see for example Malumbres-Olarte *et al*., 2017).

To reach the optimal combination of methods for any sampling effort we propose to minimize the bias of sampled diversity (TD, PD or FD) for any undersampling level compared with true (or often estimated) diversity as revealed by intensive sampling, for both inventorying and monitoring. This requires information on (near-)complete inventories for multiple sites using multiple methods. Once these data is available for any taxon, habitat or region, the following steps are:

1. Calculate fixed and variable costs per method and sample;
2. Start with a pool of samples per method. This is often 0 for all samples (an empty pool), but one might want to build on previous sampling protocols;
3. Create all possible combinations of samples per method that include the pool;
4. Calculate the cost of each combination;
5. Per combination, repeat multiple times (we used and recommend at least 1000):

a. Without replacement, randomly simulate a sampling event with the defined number of samples per method.
b. Calculate the absolute difference between sampled and true (or estimated) diversity (i.e. bias);
6. Per cost value choose the combination with the least bias.

One might also want to create a sequence of nested combinations, this way allowing for flexibility depending on the level of completeness required and resources available. In this case, in step 3 only combinations that add one sample per method are chosen and the new pool will be made up by the samples already chosen plus one sample of the method that provides the steepest slope to the accumulation curve (i.e. highest gain per cost unit).

### Extending to multiple dimensions and sites

We propose to optimize the sampling of PD and FD in addition to TD. Many ways exist to quantify PD (Pavoine & Bonsall 2011) and FD (Mammola et al. 2021), each with advantages and disadvantages, beyond the scope of this work. The principle is the same and intended to minimize the differences between sampled and true (or estimated) diversity, almost invariably taxonomic, phylogenetic or functional richness.

In addition, one should not base any sampling protocol on the experience of a single site or season. To make results comparable between these, we propose to standardize richness values between 0 and the true (or estimated) richness of each event, this way all values across sites or seasons vary between 0 and 1. This allows using multiple sampling events in a single analysis where all events have equal weight for the optimal protocol.

### Optimization of monitoring protocols (β-sampling)

While the basic algorithm was described for inventorying, hence comparisons were made for richness, i.e., α-diversity, the same principle can be applied for monitoring, i.e., detecting changes in space and time using a β-diversity framework (Tuomisto et al., 2011). Optimal monitoring requires minimizing the difference between sampled and true (or estimated) β-diversity. To guarantee such a goal, one can use any β-diversity measure of interest. We propose to use a framework that disentangles the two antithetic processes that drive overall β-diversity: species replacement and species richness differences (Carvalho et al., 2012). This framework was further expanded to PD and FD (Cardoso et al., 2014) and can be represented as:

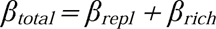

where β_total_ represents the total community variation, β_repl_ gives its fraction resulting from species replacement, i.e., the substitution of species across sites or over time, and β_rich_ accounts for the variation as a result of species richness differences, determined by the net loss / gain of species.

These measures of beta diversity are particularly robust to biases due to undersampling (Cardoso et al., 2009) and therefore the sampling effort necessary to attain low bias is small compared with α-sampling. It is also a necessary property of any monitoring protocol to be easily replicated in time, so the overall number of samples could never be as high as for inventory.

The target for the algorithm is to minimize the summed absolute difference between sampled and true (or estimated) β-diversity values using all three measures for TD, PD and FD. To make results fully comparable with alpha diversity, we propose to measure the efficiency of a combination of samples as 1 - bias, where bias is the mentioned summed difference.

### Case-studies

In the present work, we use Iberian spiders, Amazonian bats and Atlantic Forest mammals as case-studies. The ecological differences among these groups evidence that the same optimization procedure could be applied to any taxon that requires a combination of sampling methods to be fully represented, from plants and fungi to mammals. For Iberian spiders, which are a highly speciose group, we considered that an inventory should reach at least 50% of the estimated diversity. For Amazonian bats and Atlantic Forest mammals, we considered that a successful inventory should reach at least 90% of the projected diversity. For monitoring, we considered that bias should be lower than 10% in all cases. These values can be adapted to any case and are only for demonstration purposes.

### Iberian Spiders

Besides high global diversity with over 52000 known spider species (World Spider Catalog, 2024), local diversity can also be extremely high, with several hundreds of species co-existing in areas as small as 1 ha, especially in high-diversity biomes such as tropical forests (Coddington et al., 2009).

Our team sampled Iberian spiders in 18 sites using a common protocol since 2004 (Appendix 1, Table S1). In two forest sites, Arrábida and Gerês, sampled in 2004 and 2005 respectively, we collected a much larger number of samples (320) than in all others (24) and these were used previously to define the current inventory protocol for TD (COBRA) (Cardoso et al., 2008a, b; Cardoso, 2009). The remaining 16 sites were sampled between 2013 and 2014 mostly using 24 samples only, intended to capture 50% of the estimated species richness (Crespo et al., 2018). Numerous habitat types were sampled using a combination of methods: aerial searching, beating trees and branches, ground searching, sweeping and pitfall trapping (for details of the sampling methods see Cardoso et al., 2008a, b). All samples (except pitfall traps) comprised one-hour of active sampling, measured with a stopwatch and were performed both during days and nights. Diurnal and nocturnal samples of each method were considered different methods as assemblages assessed are considerably different. Pitfall traps (48) were pooled in groups of four as we estimated that four traps take an average of one hour to set up, so that they were comparable with other methods in terms of sampling effort. In this case, for simplicity and as the first demonstration of the method, we considered that there were no fixed costs, and variable costs were one per sample (hour) for every method. We also did a nested optimization, where a single sample was added to the pool at each step, according to the method that was found to be most efficient.

### Amazonian bats

Mist nets and acoustic recorders are the most used and efficient methods for sampling bats (Appel et al., 2021). However, each of these methods is biased toward different families within Chiroptera; Phyllostomidae bats being more captured by mist nets and aerial insectivorous bats (e.g., Vespertilinidae and Molossidae) being more recorded using acoustic recorders (see Appel et al., 2021; Carvalho et al., 2023). Therefore, to inventory or monitor the order Chiroptera, these two methods are considered complementary (e.g., MacSwiney et al., 2008; Carvalho et al., 2023) and we used both in a landscape composed of savannahs and forest patches in Northeastern Amazon. Each of the 12 sampled sites were surveyed for 2 nights during the rainy season. In each survey night, we installed 18 mist-nets which remained open for six hours from 15 minutes before sunset, being nine mist-nets in the savannas and nine in the forest patches, making two 110 m transects. In each of these transects, we installed three Audimoth acoustic recorders, which were set to record continuously for six hours from 15 minutes before sunset, with a sampling rate of 384 kHz and medium gain (see Carvalho et al., 2023 for details).

In this case study, we considered the costs of implementing these methods. The 18 mist-nets used in the study cost $360 each, and the aluminum poles used to install the mist-nets cost $160 each. Therefore, mist-net surveys have a fixed cost of $520, i.e. the cost of applying this method independently of the number of survey nights. However, mist-netting also has a variable cost, related to the number of nights in which the method is implemented. For each mist-net survey night we needed four researchers in the field, which represented a cost of $192 per night (i.e. $48 per researcher). Surveying a single site costs $904 ($520 + 2 nights x $192) and each additional site included in the sampling costs another $384.

For surveying with acoustic records, we have a fixed cost of $883.20, given that we used six Audiomoth acoustic recorders ($60, each), six 32 Gb storage cards ($67.20, each), and one 4 Tb hard disk to store the records ($120). The variable cost for surveying each site with acoustic records is $1062: $6 for plastic bags to protect the recorders, $192 for per-diem of two researchers to install and retrieve the recorders, and $864 of salary of one researcher to analyze and identify the acoustic records.

To evaluate the optimal protocol for α-sampling, we considered the number of sites to survey with each method. Because β-sampling involves the comparison of a number of sites, to evaluate the optimal protocol for β-sampling, we considered whether we should use mist-nets or acoustic recorders in all sites altogether.

### Atlantic forest mammals

Large and medium-sized mammals are usually surveyed through camera traps and searches in transects for direct or indirect (e.g. footprints, scats, burrows, etc.) detections, and these methods appear to be highly complementary (see Espartosa et al., 2011; Carvalho et al., 2016). We surveyed these organisms in ten transects in the Serra do Japi Biological Reserve, a 2,071 hectares protected area in Southeastern Brazil (São Paulo state) composed mostly by semi-deciduous Atlantic Forest. We installed one camera trap in each of the ten transects, and the cameras were active continuously for 45 days. We also walked these transects at an average speed of 2.7 km/h in search for direct and indirect detections of mammals. The transects were walked by two pairs of researchers, who were able to walk all the ten transects within a day. Each transect was surveyed for 40 days (see Carvalho et al., 2016 for a complete description of the methods).

Thus, we may ask whether we should use both the camera traps and the transect walks and the number of transects to include when applying these methods. In this case, we considered the per-diem payment for two researchers for two days to install and retrieve the cameras ($192 in total, $48 per researcher per day) as a fixed cost, since this value should be the same whether the researchers install one or ten cameras. Each camera trap costs $286.81, including import taxes to Brazil. For each camera, we used one 32 Gb storage card ($67.20) and 6 AA alkaline batteries ($5.50). Therefore, the variable cost of including each camera trap in the survey is $359.51. Regarding the transect walks, the fixed cost is $384, which corresponds to the per-diem payment for four researchers for two field days. In this case, there is no variable cost for transect walks.

Similarly to the Amazonian bats case-study, we evaluated the optimal protocol for α-sampling considering the number of sites to survey with each method, and for β-sampling, we considered whether we should survey with camera traps or transect walks in all sites altogether.

### Phylogenetic and Functional Diversity

PD and FD were measured as the sum of the length of branches on a phylogenetic or functional tree, respectively (Faith, 1992; Petchey & Gaston, 2002; Cardoso et al. 2014, 2024). Since TD can also be represented by a tree with each taxon linked directly to the root by a branch of unit length (star tree), tree diagrams provide a common basis for sampling optimization of TD, PD and FD (Cardoso et al., 2014, 2015, 2024).

For Iberian spiders, a phylogenetic tree was built using the procedure detailed in Macías-Hernández et al. (2020a). In short, a concatenate matrix of the mitochondrial cytochrome c oxidase subunit I (COI) and the nuclear 28S rRNA (28S) genes sequenced for the 372 species collected in the 16 sites was constructed (see further details on laboratory protocols and molecular data in Macías-Hernández et al., 2020a). Approximately 1000 additional species representing most spider families, sampled for the same two gene regions, together with four additional genes (12S rRNA,16S rRNA, histone H3 and 18S rRNA), available from a previous study (Wheeler et al., 2016), were added to the COI and 28S concatenate matrix. Then, a Maximum Likelihood (ML) phylogenetic tree was inferred using the program RAxML-HPC v. 8.2.12 (Stamatakis, 2014), remotely run on XSEDE at the CIPRES Science Gateway (Miller et al., 2010). The phylogenetic analysis was performed with a topological constraint provided by a phylotranscriptomic study on spiders (Fernández et al., 2018), which also provided the calibration points to infer time-stamped phylogeny. The resulting ML tree was made ultrametric using the *congruify.phylo* function in the R package *GEIGER* (Harmon et al., 2008), and the resulting table of calibration points was used to date the target trees using the program treePL (Smith & O’Meara, 2012). This tree was used for the optimization of monitoring PD.

In lieu of precise phylogenetic information on the member taxa of a community, PD can also be calculated from an ultrametric tree representing the Linnaean hierarchy (Ricotta et al., 2012). As no genetic data was available for the two sites used for the optimization of inventory, a tree based on taxonomic hierarchy was constructed as a surrogate using the function *linnean* in the R package *BAT* (Cardoso et al. 2015). This tree was used for the optimization of inventorying PD.

For Amazonian bats and Atlantic Forest mammals, we obtained an ultrametric consensus tree from the 1000 trees retrieved from Upham et al. (2019) using the R package *ape* (Paradis & Schliep, 2019). This phylogeny was based on a 31-gene supermatrix and included all the 48 and 28 species detected in the study, respectively.

For Iberian spiders, we constructed a functional tree using data from 11 functional traits, namely: mean total body size (males and females), prosoma length, prosoma width, prosoma height, tibia I length, fang length, dispersal ability (ballooning propensity: (F) frequent; (O) occasional or (R) rare), vertical stratification (from epigean to arboreal), circadian activity (nocturnal / diurnal), foraging strategy (no web / web builders: capture web, sensing web, tube web, sheet web, space web and orb web), and trophic specialization (specialist or generalist), based on bibliographic searches, personal knowledge of species or derived from our own sampling data (further details in Cardoso et al., 2011b, Macías-Hernández et al., 2020b). To avoid correlation between prosoma width and other morphological variables, we calculated the residual values of each morphological variable against prosoma width. A functional dissimilarity matrix between species was constructed using Gower’s distance and rescaling variables by selecting specific weights. Each morphological variable (but prosoma width) and the binary variables related to types of webs were weighted as 1/n types. The functional tree was built as a neighbor-joining tree using the function *tree.build* of the R package *BAT* (Cardoso et al., 2015). Using neighbor-joining is the only representation that allows comparability with analyses of PD using phylogenetic trees, an unbiased representation of species distances, and to use the beta diversity partitioning framework for the optimization of the monitoring protocol (Cardoso et al., 2024).

For Amazonian bats, we constructed a functional tree based on four traits: logarithmized body mass, forearm size, trophic level (phytophagous or animalivorous), and dietary guild (frugivore, insectivore, aerial insectivore, omnivore or sanguinivore). Body mass and forearm size were obtained from individuals captured in the field, and the trophic guild was based on Giannini and Kalko (2004). Diet was based on a repository of published papers that analyzed the diet of different species of bats around the world (Alroy, 2017). A functional tree was built the same way as for spiders.

For Atlantic Forest mammals we used the following eight traits: logarithmized body mass, vertical strata (terrestrial, scansorial or arboreal), three binary traits indicating whether the species presents diurnal, nocturnal, and/or crepuscular activity, and three traits corresponding to the percentage of animal prey, reproductive plant parts, and vegetative plant parts in the species’ diet. These traits were obtained from Wilman et al. (2014). To account for the correlations between some of these traits, we used the Gower’s distance on weighted traits to balance the contribution of each trait to the dissimilarity matrix using the *gawdis* function (de Bello et al. 2021). We assigned the three activity periods and the three diet traits of the Atlantic Forest mammals as groups in this analysis. Then, we used the *tree.build* function in *BAT* to build the functional trees again using neighbor-joining.

For all taxa, we used the functions *optim.alpha* and *optim.beta* in *BAT* (Cardoso et al., 2015) to find the best allocation of sampling effort for inventorying and monitoring, respectively.

## Results

### Iberian spiders

A total of 272 species and 13,064 adult individuals (from two sites) were used for optimizing α-sampling, and 587 species and 18,623 adults (from 16 sites) for optimizing β-sampling. As we previously found that six hours is the maximum effort a single researcher can perform per day without fatigue influencing results (Cardoso 2009), we looked for combinations that would add to a total number of samples multiple of six. For α-diversity, the optimal combination of 24 samples for both TD and PD was very similar (Table 2). In both cases, the proportion of diversity sampled was higher than 50%, reaching 57% for PD. The optimal protocol was close or even over the upper confidence limit for random sampling for most of the accumulation process (Fig. 1). For FD, the main difference laid on the optimality of doing ground searching during the day instead of beating or sweeping. In any case, as many species are very similar in their traits, the sampled diversity with just 24 samples was very high, reaching 79% of true FD. This was confirmed by the much faster accumulation rate of FD with increasing number of samples (Fig. 1). In the constrained exercise, the algorithm was able to rapidly reach method combinations and sampling levels on par with the unconstrained optimization (Table 2; Fig. 1).

**Fig 1.**
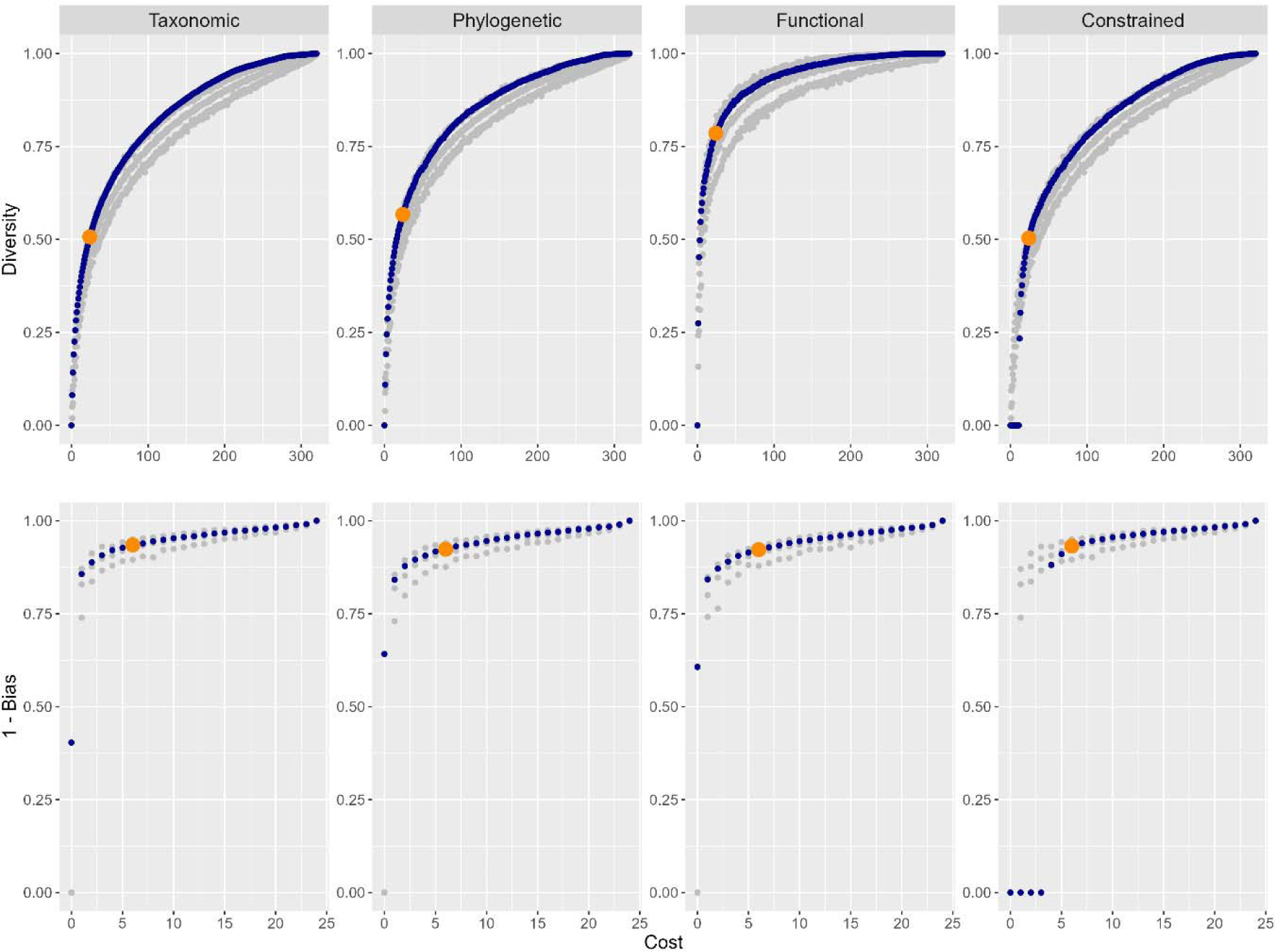
Spider sampling optimization. Accumulation curves for optimized inventorying (α-sampling; upper panels) and monitoring (β-sampling; lower panels) of taxonomic, phylogenetic, and functional diversity of spiders, plus the optimization of taxonomic diversity constrained to an initial sample of 12 and four pitfall traps, respectively, for α and β-sampling (blue dots). The results for β-sampling are presented as 1 - bias, so, the greater the value, the higher the accuracy. The orange dots represent the optimal sampling protocols, i.e. with 24 and six samples, respectively (see text for details). Grey dots represent the median and 95% confidence limits of a random choice of methods along the accumulation curve.

**Table 2.**
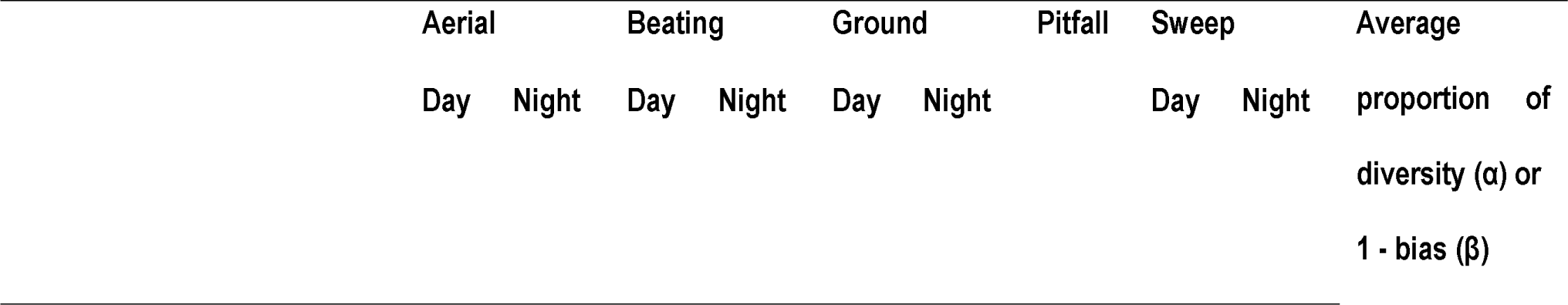
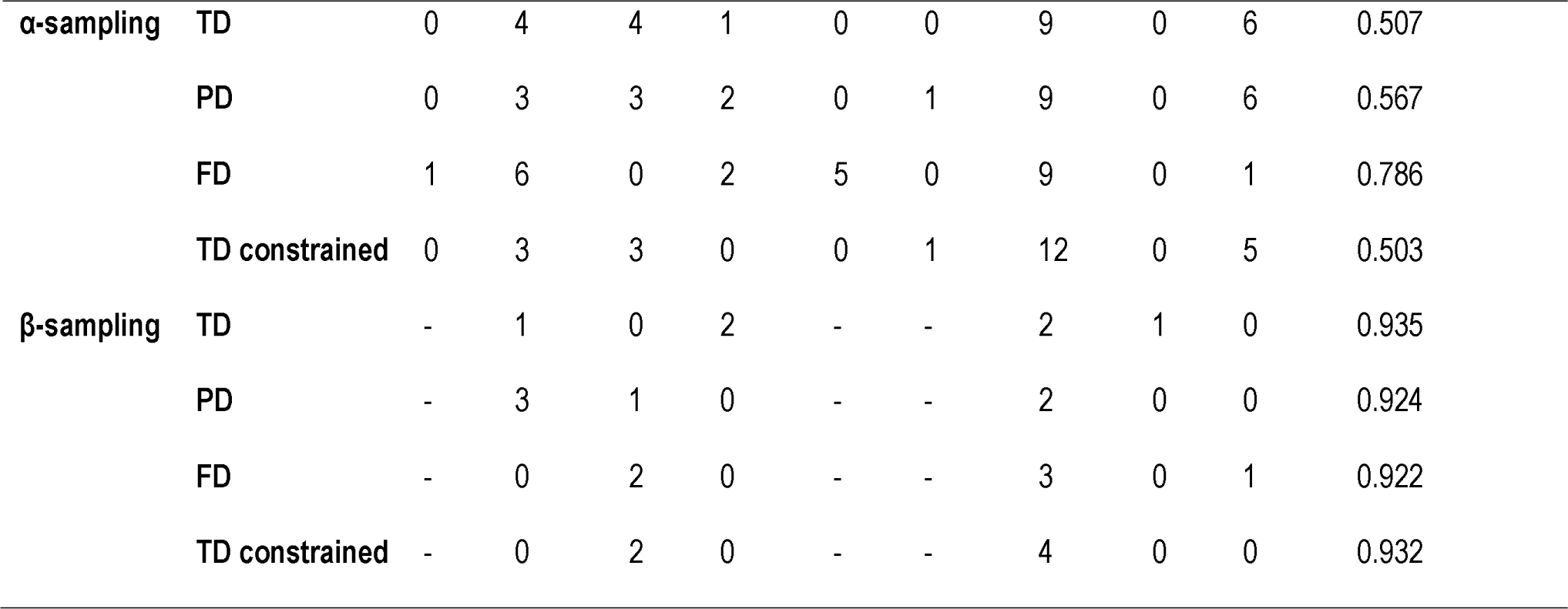
Spider sampling optimization.

Optimal number of samples per method for inventorying (α-sampling) and monitoring (β-sampling) of spider taxonomic (TD), phylogenetic (PD) and functional (FD) diversity. A total of 24 and six samples were chosen respectively (see text for details). The constrained runs start with 12 or four pitfall trap samples, respectively.

For β-diversity, the optimal combination of six samples for all measures was similar (Table 2). In all cases, the 1 - bias from true diversity was between 0.922 (for FD) and 0.935 (for TD). The optimal protocol was however not significantly different from a random one based on the inventory protocol (Fig. 1). In the constrained exercise, the algorithm was able to rapidly reach method combinations and sampling levels on par with the unconstrained optimization (Table 2; Fig. 1).

### Amazonian bats

We had 234 records from 48 bat species (155 records and 38 species in mist-nets and 79 records and 12 species in acoustic recorders). Despite the higher costs and lower diversity sampled by acoustic recorders, they recorded 10 unique species. Therefore, for taxonomic α-diversity, the most cost-effective combination for inventorying bats is using mist-nets in 12 sites and acoustic recorders in a single site (Fig. 2). This combination returned 89.5% of the diversity with 37.7% of the cost of the full inventorying scheme. For phylogenetic α-diversity, the choice of the best solution is more subjective, given that there is a more gradual increase of the PD with the cost of implementing the methods. For example, using 5 mist-net sites and 1 acoustic recorder site results in 84.5% of the PD with 23.4% of the costs of the full scheme. Adding more mist-net sites (i.e. 10 sites) while still using a single acoustic recorder increases the recorded PD to 90.3%, but also increases the costs to 33.4% of the full scheme. With 12 mist-net sites and three acoustic recorders, we achieve 94.8% of the PD with 49.0% of the cost of the full scheme (Fig. 2). Therefore, the decision considering this trade-off between higher diversity and higher costs must be made taking into account the resources available for the study. Similarly to the TD, for functional α-diversity, the most cost-effective solution is surveying in 12 mist-netting sites and no acoustic recording sites, which returned 94.5% of the FD with 27.3% of the costs of the full scheme (Fig. 2).

**Fig 2.**
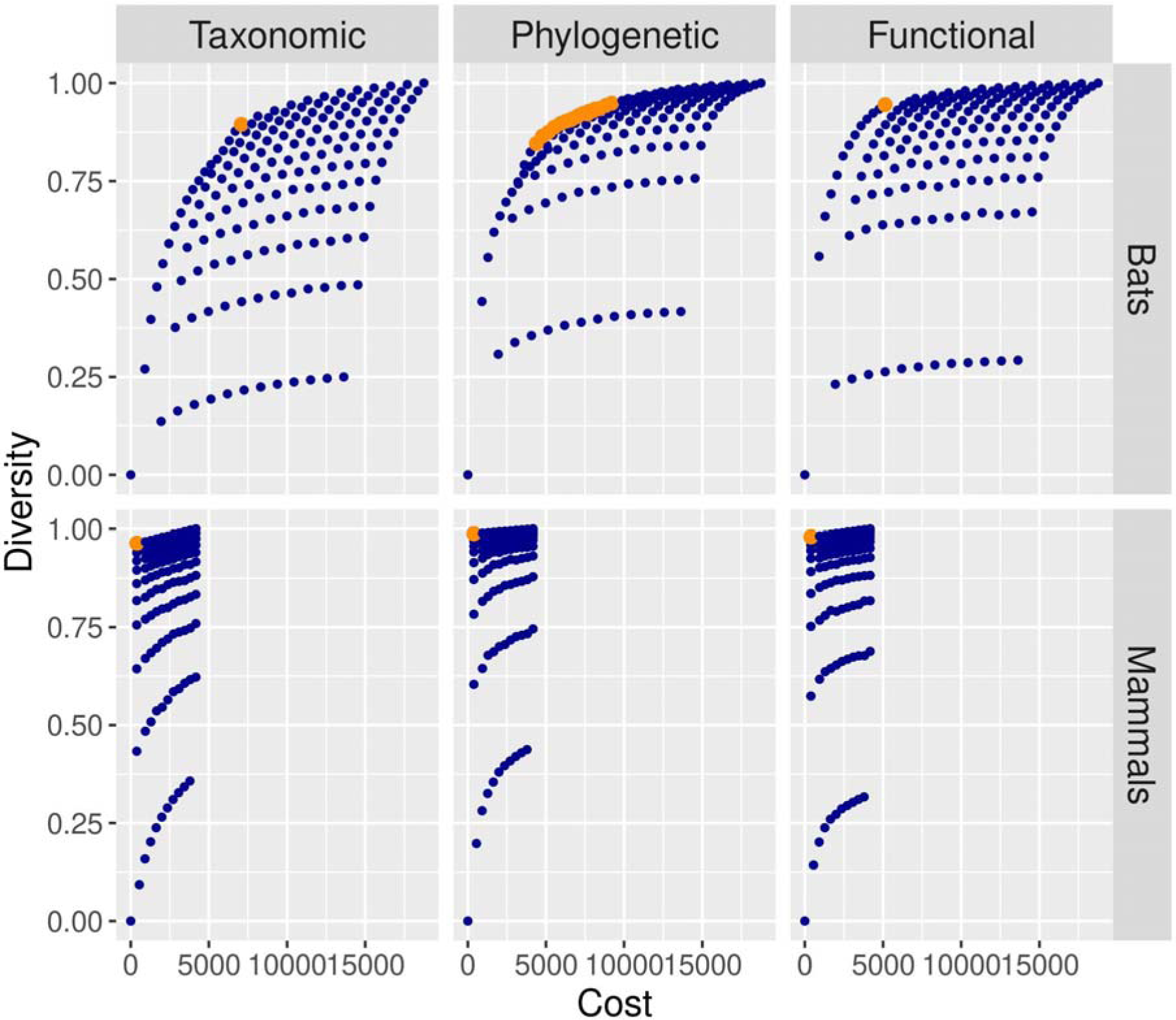
Mammal inventorying optimization. Relationship between the costs of implementing the inventorying scheme and the resulting α-diversity. The results are shown for taxonomic, phylogenetic and functional diversity of Amazonian bats and Atlantic Forest mammals. α-diversity is scaled from 0 (no diversity) to 1 (diversity recorded in the scheme containing all the methods). The dots represent all the possible combinations of methods used to survey bats and mammals in the case-studies. In the bat case study, we used mist-nets and acoustic recorders. In the mammal study case, we used camera traps and transect walks. The most cost-effective schemes are in orange.

For biodiversity monitoring, the best sampling scheme is to sample only with mist-nets for TD, PD, and FD, which results in biases <10% (Fig. 3).

**Fig 3.**
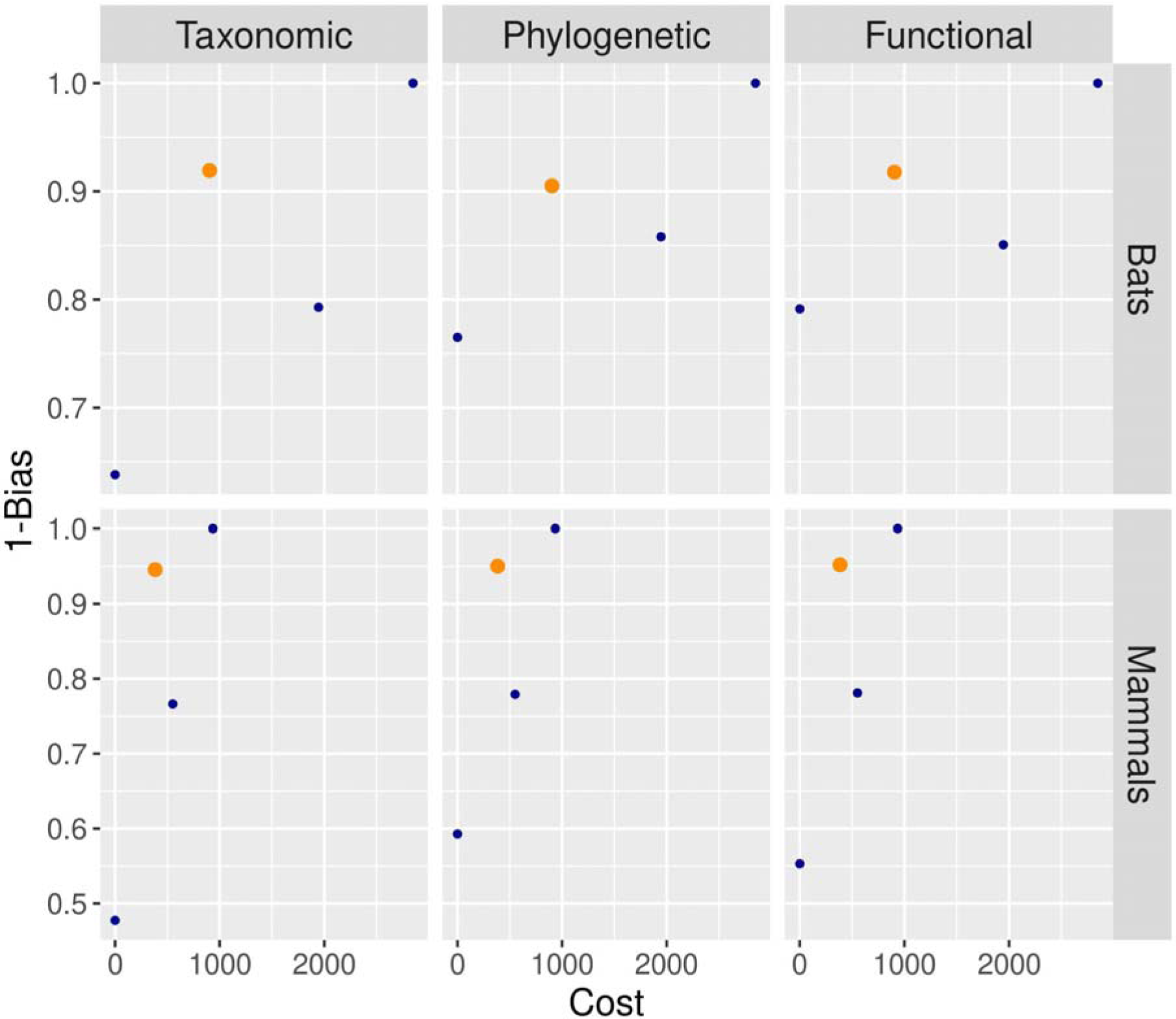
Mammal monitoring optimization. Relationship between the costs of implementing the monitoring scheme and the resulting bias in the assessment of the β-diversity. The results are shown for taxonomic, phylogenetic and functional diversity of Amazonian bats and Atlantic Forest mammals. The results are presented as 1 - bias, so, the greater the value, the lower the bias. The dots represent all the possible combinations of methods used to survey bats and mammals in the case-studies. In the bat case study, we used mist-nets and acoustic recorders. In the mammal study case, we used camera traps and transect walks. The most cost-effective schemes are in orange.

### Atlantic Forest mammals

We had 439 records of mammals from 28 species (28 records and 10 species in camera traps and 391 records and 27 species in transect walks). Considering the higher cost and lower detectability of camera traps, the combination of samples which resulted in the highest α-diversity with lower cost is sampling using 10 transects and no camera traps. The results were similar for TD, PD and FD (Fig. 2). For biodiversity monitoring, the results are similar and sampling only in transects results in a relatively low bias (∼5%) with a third of the costs (Fig. 3).

## Discussion

Here we propose a method to optimize and standardize the inventory and monitoring of taxonomic, phylogenetic and functional diversity, and further introduce the definitions of α-sampling and β-sampling. We demonstrate the method using Iberian spiders, Amazonian bats, and Atlantic Forest mammals as case-studies. This same method is applicable to any taxon at any region, as long as sampling can be done using a combination of methods. In fact, a first test for monitoring of TD was already proposed for Macaronesian spiders, beetles and bryophytes (Borges et al., 2018), proving the usefulness of the concept for different taxa.

Instead of relying on subjective opinions of researchers, this approach allows the proposed protocols to meet six requirements that facilitate their widespread use: (1) the protocols are suitable for the purpose and target organisms, (2) they are efficient by maximizing the gain/effort ratio, (3) feasible by requiring minimum resources, (4) flexible as different sampling efforts can be tested and proposed, (5) transparent as the process to reach them is explicit, and (6) accountable as results can be tested *a posteriori* (Cardoso, 2009).

The algorithm proposed here can support researchers and environmental managers to choose optimal sampling schemes for inventory and monitoring. However, the choice of the optimal sampling scheme may depend on the availability of money, time, and human resources. Although here we only considered money as a cost parameter to support the choice of the best sampling scheme, the demand for time or human resources can also be included as cost parameters in the algorithms. The choice of the most adequate cost parameter depends on which factor is more limiting to the inventory/monitoring project. In some cases, the researchers or environmental managers may even choose sub-optimal sampling schemes if the resource availability is very limited. In this case, the proposed algorithm is useful to indicate how much of the diversity is being assessed in the inventory or the amount of bias that is being introduced in the monitoring.

We chose to focus on trees as representations for both PD and FD because they allow the comparison of TD, PD and FD under a common framework (Cardoso et al., 2014, 2015, 2024). This optimization method can however be adapted to other representations, such as multidimensional kernel-density hypervolumes, a common representation for FD (Blonder et al., 2018; Carvalho & Cardoso, 2020; Mammola & Cardoso, 2020; Mammola et al. 2021).

For Iberian spiders, the monitoring protocol we propose was delineated using methods targeting different micro-habitats (Table 2). This probably explains why a combination of several different methods is optimal, as different components of the assemblages should be targeted. Different habitats could require a different combination of methods, depending on, among others, existence of an arboreal stratum. Optimizing β-sampling may therefore depend on the goals and should be done for each biome, region and/or habitat independently. This is further reinforced by the fact that β-diversity is usually unitless, so it is possible to compare protocols using different methods and efforts. Contrary to inventorying, which can be, and is being done, at a global level using very similar methodology, allowing comparisons at large to global scales, monitoring is case-specific, and results are often context-dependent.

For Amazonian bats, the optimal sampling protocol varied between TD, PD, and FD for α-sampling. Therefore, if a project aims to use different diversity types, it is important to assess the best options for all the diversity types to avoid choosing sub-optimal protocols. Nevertheless, ideally, before starting any kind of monitoring of a site or region, an inventory should be conducted, so that a baseline or control can be established (Stork et al., 1996). Inventories provide the baseline that allows perceiving if changes in community composition are due to changes in the abundances of species, clades or traits already living but previously rare in the site, or to the replacement (i.e. β_repl_), extirpation or arrival (i.e. β_rich_) of new components to the community (Carvalho et al., 2012).

Uncoupling of real vs. sampling artifact related change in monitoring protocols is a hard task and deserves careful consideration. First, the data used to define the protocol is of critical importance. It must be extensive, from multiple sites covering the entire spectrum we want to monitor, to provide as good a baseline as possible. Second, we must be careful choosing the β-diversity measures used to optimize the protocol. The ones used here are much less prone to error due to undersampling than either α-diversity or other β-diversity measures (Cardoso et al., 2009). Also, careful consideration should be given to the details of implementation of the methods. For example, in the case-study of Iberian spiders we always sampled 100×100m plots and these protocols are only comparable within similar sampled areas. In the case study of Atlantic Forest mammals, the cameras were deployed for 45 days, and leaving the cameras for longer periods in the field may change the optimal sampling scheme. In the case study of Amazonian bats, the choice of the best sampling scheme should depend on the number of mist-nets used in each sampling site.

Finally, one may combine simultaneous α and β-sampling. One may α-sample several reference sites and β-sample additional sites with the sole purpose of comparing them with the reference condition. This might be useful for example to test change in community composition along a disturbance gradient, test the effect of habitat margins, or study meta-population source-sink dynamics in a fragmented landscape.

### Setting up a sampling scheme

A few considerations must be made before the replication of any sampling protocol. The spatial extent and temporal scale must be determined in view of specific objectives and available resources. The number and location of sampling sites could, as much as possible, be consistent with some procedure common for an entire scheme. Many projects may benefit from maximizing environmental diversity covered by sampling (including habitat types), with the reasoning that environmental and biological diversity may be correlated (Faith & Walker, 1996; Hortal & Lobo, 2005). This allows maximizing the overall diversity captured and, most importantly, the range of variation in biodiversity of a region. One must remember though, that if monitoring is to be done, the sites should be fixed, and ease of accessibility may be important.

For monitoring, the frequency and duration of the scheme is crucial. Often it is not possible to detect changes in species composition or population abundance before a relatively large time lag (e.g., Wauchope et al. 2019; White, 2019). It is also important to be able to tell apart consistent change and natural fluctuations. Thus, besides specimen sampling, it is important to collect the most relevant biotic and abiotic variables. Usually habitat type, temperature, precipitation, tree and ground cover during or immediately before the time of sampling could potentially be useful to help understanding community composition and change. Precipitation is known to strongly influence the activity of many animals and collectors should therefore make an effort to only sample in appropriate conditions (Burner et al., 2020).

In many cases, voucher specimens, or better yet entire collections, should be preserved as only this way it is possible to guarantee future comparability of projects if taxonomy changes or if further analyses are needed relying on genetic or functional data. One may for example want to sequence new genes that allow the comparison of intra-specific genetic diversity of given species between sites or years even if these comparisons were not thought of from the beginning of the project. One may also want to detect changes in functional traits such as body size driven by environmental change. All projects involving specimen sampling should therefore plan and allocate appropriate funding to specimen deposition in public collections such as in natural history museums (Schilthuizen et al., 2015).

### Conclusions

The widespread use of similar sampling protocols at global levels and regular repetition in time can potentially have a major impact on the scope and consequently usefulness of projects such as GEO-BON, RLI and LPI. But the impact goes much beyond these. Often researchers use sampling methods and effort per method with which they are more familiar with or that are specific to a given project, necessarily limited in space and time. Besides potentially low cost-effectiveness, this creates data that are not comparable with other datasets collected by different or even the same researchers for other projects. The comparable sampling of biodiversity in space and/or time allows reusing of data collected for specific purposes, potentiating a synergistic effect among different projects. This makes data useful much beyond their initial plan. We therefore strongly advocate the optimization, standardization and widespread adoption of sampling protocols for all taxa at a global level, for both inventory and monitoring of all levels of biodiversity: taxonomic, phylogenetic and functional.

## Acknowledgments

We thank all the collectors of past and present studies that contributed to the dataset used in this paper: Luis Carlos Crespo, Rui Carvalho, Alberto de Castro, Cristina Frías-López, Pedro Humberto Castro, Clara Gaspar, Ana Filipa Gouveia, Sérgio Henriques, Eva de Mas, Paola Mazzuca, Elisa Mora, Jordi Moya, Vera Opatova, Luis Carlos Pereira, Enric Planas, Carles Ribera, Marcos Roca-Cusachs, Dolores Ruiz, Nikolaj Scharff, Jesper Schmidt, Israel Silva, Ricardo Silva, Pedro Sousa, Tamás Szüts, Vanina Tonzo, João D. Miguel, Adrià LópezLJBaucells, Ricardo Rocha, Jorge M. Palmeirim, Isaí Castro, Bruna S. Xavier, Karen Mustin, Daniela Rato, Cledinaldo Marques, Cremilson Marques, Gustavo Silveira, Joandro Pandilha, Jackson Souza, Luís Miguel Rosalino, Maíra S. M. Godoy, Renan M. Dias, Carlos E. L. Esbérard and Cristina H. Adania. MA was supported by grant 485/2012 of the Organismo Autónomo de Parques Nacionales (Ministerio de Agricultura, Alimentación y Medio Ambiente). NMH was supported by the H2020-MSCA-IF-2015 Programme (BIODIV ISLAND-CONT project, Grant no. 706482). WDC is supported by ‘Ayudas Maria Zambrano’ (CA3/RSUE/2021-00197), funded by the Spanish Ministry of Universities.

## Authors’ contributions

PC and RH conceived the ideas, designed methodology and led the writing of the manuscript; PC, MAA, NMH and WC collected the data; PC, MAA, NMH, JCC and RH analysed the data. All authors contributed critically to the drafts and gave final approval for publication.

## Data availability

All code to replicate the analyses is deposited at https://github.com/cardosopmb/optimSampling

Spider distribution data is fully available in https://doi.org/10.3897/BDJ.6.e29443

Genetic data is fully available in https://www.mdpi.com/1424-2818/12/8/288/s1

Functional data is fully available in https://doi.org/10.6084/m9.figshare.8320004.v3

Bat diversity data is fully available in https://doi.org/10.1007/s42974-022-00131-5

Mammal diversity data is fully available in https://doi.org/10.1590/1678-4766e2016005 and https://seer.ufu.br/index.php/biosciencejournal/article/view/14258.

## Appendix S1

**Table S1.**
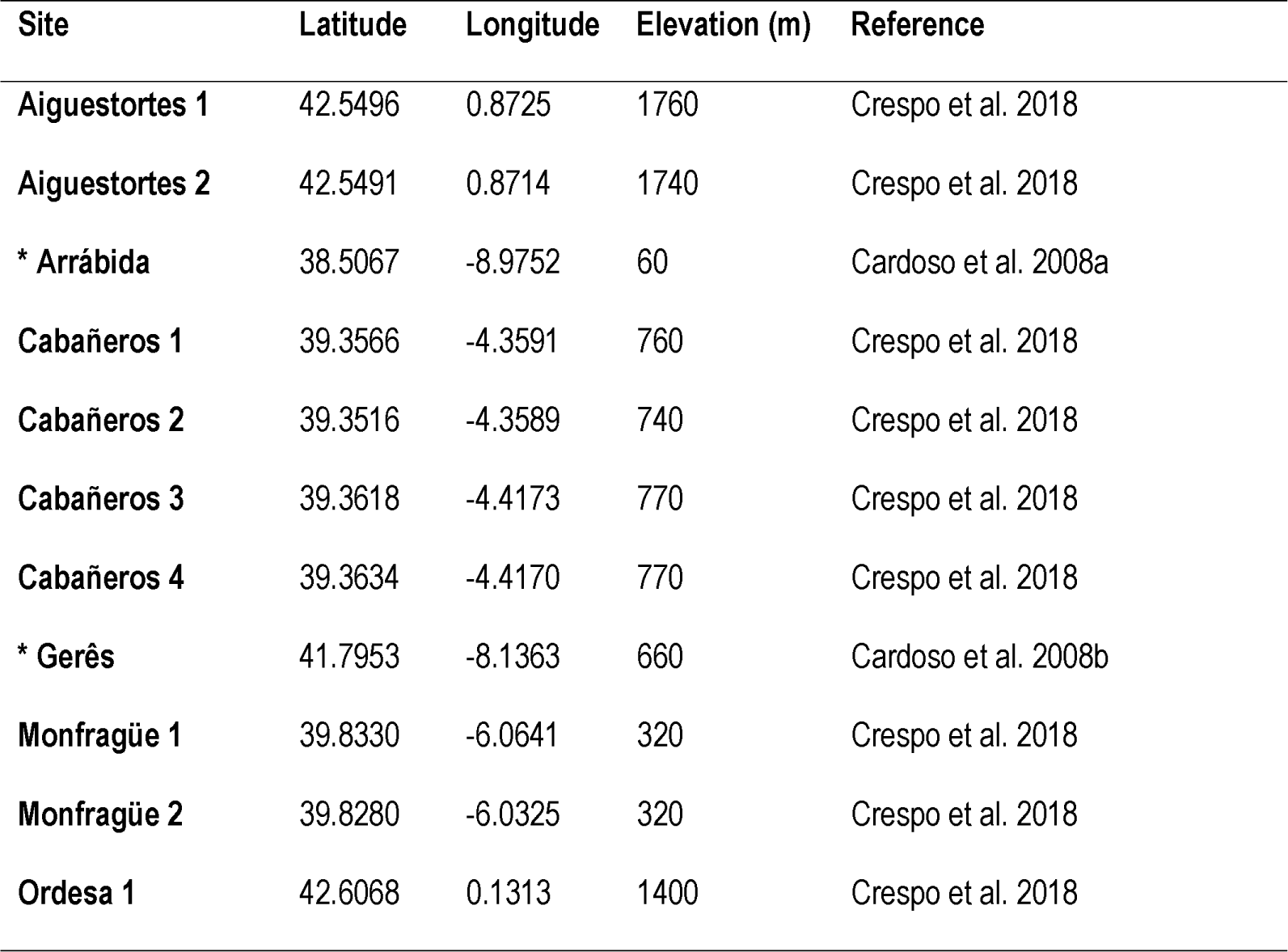
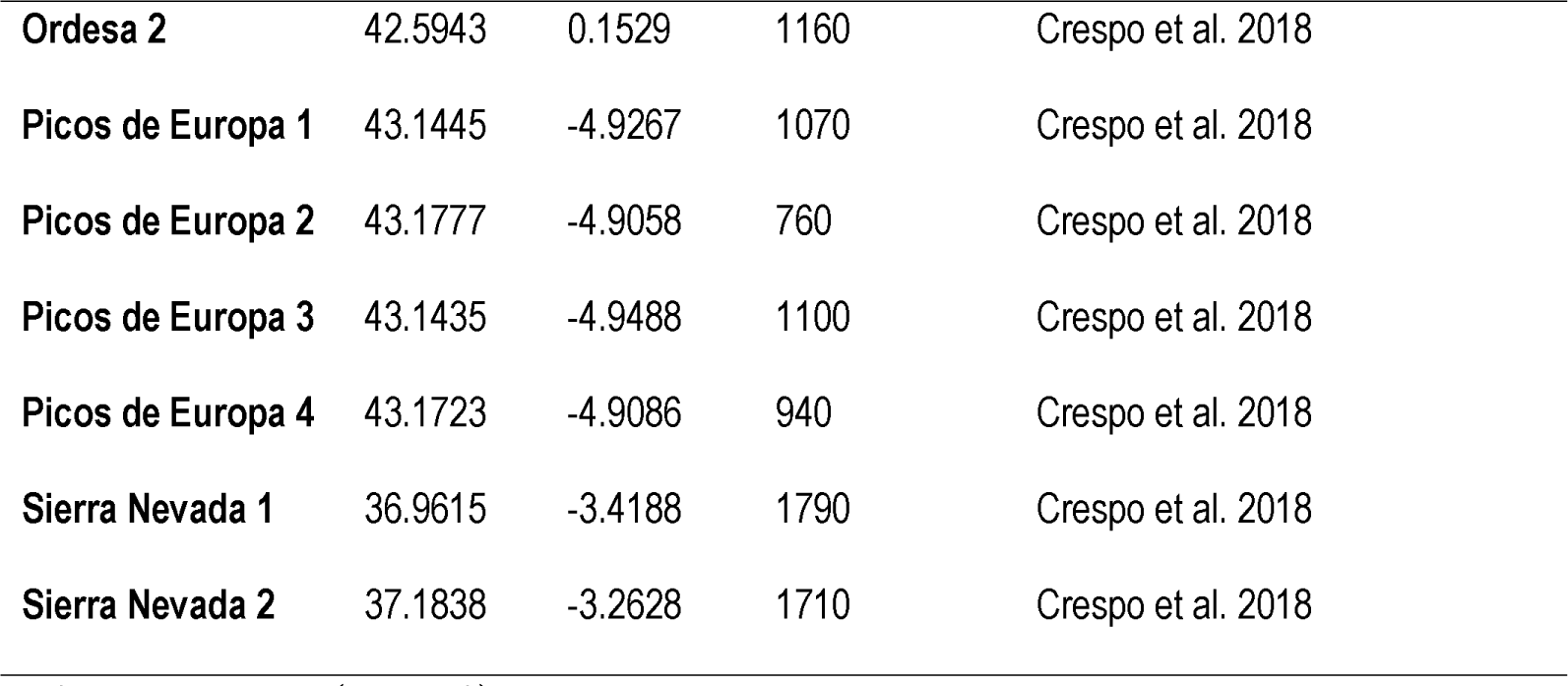
Eighteen sampled sites in Iberian oak forests.

## References

Alroy J. Effects of habitat disturbance on tropical forest biodiversity. Proc Natl Acad Sci U S A. 2017;114(23):6056–6061.

Appel G, Capaverde UD Jr, de Oliveira LQ, Pereira LGA, Tavares VC, López-Baucells A, et al. Use of complementary methods to sample bats in the Amazon. Acta Chiropterologica. 2021;23:499–511.

Bawa KS, Menon S. Biodiversity monitoring: the missing ingredients. Trends Ecol Evol. 1997;12:42–43.

Blonder B, Morrow CB, Maitner B, Harris DJ, Lamanna C, Violle C, et al. New approaches for delineating n-dimensional hypervolumes. Methods Ecol Evol. 2018;9:305–319.

Borges PAV, Cardoso P, Kreft H, Whittaker RJ, Fattorini S, Emerson BC, et al. A Global Island Monitoring Scheme (GIMS) for the long-term coordinated survey and monitoring of forest biota across islands. Biodivers Conserv. 2018;27:2567–2586.

Burner RC, Birkemoe T, Olsen SL, Sverdrup-Thygeson A. Sampling beetle communities: Trap design interacts with weather and species traits to bias capture rates. Ecol Evol. 2020;10:14300–14308.

Butchart SH, Walpole M, Collen B, Van Strien A, Scharlemann JP, Almond RE, et al. Global biodiversity: indicators of recent declines. Science. 2010;328(5982):1164-1168.

Cardoso P. Standardization and optimization of arthropod inventories – the case of Iberian spiders. Biodivers Conserv. 2009;18:3949–3962.

Cardoso P, Leather S. Predicting a global insect apocalypse. Insect Conserv Divers. 2019;12:263–267.

Cardoso P, Gaspar C, Pereira LC, Silva I, Henriques SS, Silva RR, et al. Assessing spider species richness and composition in Mediterranean cork oak forests. Acta Oecologica. 2008a;33:114–127.

Cardoso P, Scharff N, Gaspar C, Henriques SS, Carvalho R, Castro PH, et al. Rapid biodiversity assessment of spiders (Araneae) using semi-quantitative sampling: a case study in a Mediterranean forest. Insect Conserv Divers. 2008b;1:71–84.

Cardoso P, Borges PAV, Veech JA. Testing the performance of beta diversity measures based on incidence data: the robustness to undersampling. Divers Distrib. 2009;15:1081–1090.

Cardoso P, Erwin TL, Borges PAV, New TR. The seven impediments in invertebrate conservation and how to overcome them. Biol Conserv. 2011a;144:2647–2655.

Cardoso P, Pekár S, Jocqué R, Coddington JA. Global patterns of guild composition and functional diversity of spiders. PLoS One. 2011b;6:e21710.

Cardoso P, Rigal F, Carvalho JC, et al. Partitioning taxon, phylogenetic and functional beta diversity into replacement and richness difference components. J Biogeogr. 2014;41:749–761.

Cardoso P, Rigal F, Carvalho JC. BAT - Biodiversity Assessment Tools, an R package for the measurement and estimation of alpha and beta taxon, phylogenetic and functional diversity. Methods Ecol Evol. 2015;6:232–236.

Cardoso P, Barton PS, Birkhofer K, Chichorro F, Deacon C, Fartmann T, et al. Scientists’ warning to humanity on insect extinctions. Biol Conserv. 2020;242:108426.

Cardoso P, Guillerme T, Mammola S, Matthews TJ, Rigal F, Graco-Roza C, et al. Calculating functional diversity metrics using neighbor-joining trees. bioRxiv. 2024. 10.1101/2022.11.27.518065.

Carvalho JC, Cardoso P. Decomposing the causes for niche differentiation between species using hypervolumes. Front Ecol Evol. 2020;8:243.

Carvalho JC, Cardoso P, Gomes P. Determining the relative roles of species replacement and species richness differences in generating beta-diversity patterns. Global Ecol Biogeogr. 2012;21:760–771.

Carvalho WD, Rosalino LM, Adania CH, Esbérard CE. Mammal inventories in Seasonal Neotropical Forests: traditional approaches still compensate drawbacks of modern technologies. Iheringia. Série Zoologia. 2016;106.

Carvalho WD, Miguel JD, da Silva Xavier B, Lopez-Baucells A, de Castro IJ, Hilario RR, et al. Complementarity between mist-netting and low-cost acoustic recorders to sample bats in Amazonian rainforests and savannahs. Community Ecol. 2023;24(1):47–60.

Coddington JA, Agnarsson I, Miller JA, Kuntner M, Hormiga G. Undersampling bias: the null hypothesis for singleton species in tropical arthropod surveys. J Anim Ecol. 2009;78:573–584.

Cowie RH, Bouchet P, Fontaine B. The Sixth Mass Extinction: fact, fiction or speculation? Biol Rev. 2022;97(2):640–663.

Crespo LC, Domènech M, Enguídanos A, Malumbres-Olarte J, Cardoso P, Moya-Laraño J, et al. A DNA barcode-assisted annotated checklist of the spider (Arachnida, Araneae) communities associated with white oak woodlands in Spanish National Parks. Biodivers Data J. 2018;6:e29443.

Danielsen F, Jensen AE, Alviola PA, Balete DS, Mendoza M, Tagtag A, et al. Does monitoring matter? A quantitative assessment of management decisions from locally-based monitoring of protected areas. Biodivers Conserv. 2005;14:2633–2652.

de Bello F, Botta-Dukát Z, Lepš J, Fibich P. Towards a more balanced combination of multiple traits when computing functional differences between species. Methods Ecol Evol. 2021;12(3):443–448.

Devictor V, Mouillot D, Meynard C, Jiguet F, Thuiller W, Mouquet N. Spatial mismatch and congruence between taxonomic, phylogenetic, and functional diversity: the need for integrative conservation strategies in a changing world. Ecol Lett. 2010;13:1030–1040.

Emerson BC, Casquet J, López H, Cardoso P, Borges PAV, Mollaret N, et al. A robust field survey protocol for comparing forest arthropod biodiversity across spatial scales. Mol Ecol Resour. 2017;17:694–707.

Espartosa KD, Pinotti BT, Pardini R. Performance of camera trapping and track counts for surveying large mammals in rainforest remnants. Biodivers Conserv. 2011;20:2815–2829.

Faith DP. Conservation evaluation and phylogenetic diversity. Biol Conserv. 1992;61:1-10.

Faith DP, Walker PA. Environmental diversity: on the best-possible use of surrogate data for assessing the relative biodiversity of sets of areas. Biodivers Conserv. 1996;5:399–415.

Fernández R, Kallal RJ, Dimitrov D, Ballesteros JA, Arnedo MA, Giribet G, et al. Phylogenomics, diversification dynamics, and comparative transcriptomics across the spider Tree of Life. Curr Biol. 2018;28:1489-1497.

Gardner TA, Barlow J, Araujo IS, Ávila-Pires TC, Bonaldo AB, Costa JE, et al. The cost-effectiveness of biodiversity surveys in tropical forests. Ecol Lett. 2008;11(2):139–150.

Giannini NP, Kalko EK. Trophic structure in a large assemblage of phyllostomid bats in Panama. Oikos. 2004;105:209–220.

Harmon LJ, Weir JT, Brock CD, Glor RE, Challenger W. GEIGER: investigating evolutionary radiations. Bioinformatics. 2008;24:129–131.

Henriques SS, Böhm M, Collen B, Luedtke J, Hoffmann M, Hilton-Taylor C, et al. Accelerating the monitoring of global biodiversity: revisiting the sampled approach to generating Red List Indices. Conserv Lett. 2020;e12703.

Hilty J, Merenlender AM. Faunal indicator taxa selection for monitoring ecosystem health. Biol Conserv. 2000;92:185–197.

Hortal J, Lobo JM. An ED-based protocol for optimal sampling of biodiversity. Biodivers Conserv. 2005;14:2913–2947.

Hortal J, de Bello F, Diniz-Filho JAF, Lewinsohn TM, Lobo JM, Ladle RJ. Seven shortfalls that beset large-scale knowledge of biodiversity. Annu Rev Ecol Evol Syst. 2015;46:523–549.

Jetz W, McGeoch MA, Guralnick R, Ferrier S, Beck J, Costello MJ, et al. Essential biodiversity variables for mapping and monitoring species populations. Nat Ecol Evol. 2019;3:539–551.

Kiljunen N, Pajunen T, Fukushima CS, Soukainen A, Kuurne J, Korhonen T, et al. Standardized spider (Arachnida, Araneae) inventory of Kilpisjärvi, Finland. Biodivers Data J. 2020;8:e50775.

Leakey RE, Lewin R. The Sixth Extinction: Biodiversity and Its Survival. Science Masters. ISBN-13: 978–0297817338; 1996.

Ledger SE, Loh J, Almond R, Böhm M, Clements CF, Currie J, et al. Past, present, and future of the Living Planet Index. npj Biodiversity. 2023;2(1):12.

Longino JT, Colwell RK. Biodiversity assessment using structured inventory: capturing the ant fauna of a tropical rain forest. Ecol Appl. 1997;7:1263–1277.

Macías-Hernández N, Domenech M, Cardoso P, Emerson B, Borges P, Lozano-Fernandez J, et al. Building-up of a robust, densely sampled spider tree of life for assessing phylogenetic diversity at the community level. Diversity. 2020a;12:288.

Macías-Hernández N, Ramos C, Domenech M, Febles S, Santos I, Arnedo MA, et al. A database of functional traits for spiders from native forests of the Iberian Peninsula and Macaronesia. Biodivers Data J. 2020b;8:e49159.

MacSwiney GMC, Clarke FM, Racey PA. What you see is not what you get: The role of ultrasonic detectors in increasing inventory completeness in Neotropical bat assemblages. J Appl Ecol. 2008;45:1364–1371.

Malumbres-Olarte J, Scharff N, Pape T, Coddington JA, Cardoso P. Gauging megadiversity with optimized and standardized sampling protocols: A case for tropical forest spiders. Ecol Evol. 2017;7:494–506.

Mammola S, Cardoso P. Functional diversity metrics using kernel density n-dimensional hypervolumes. Methods Ecol Evol. 2020;11:986–995.

Mammola S, Carmona CP, Guillerme T, Cardoso P. Concepts and applications in functional diversity. Funct Ecol. 2021;35:1869–1885.

Mammola S, Adamo M, Antić D, Calevo J, Cancellario T, Cardoso P, et al. Drivers of species knowledge across the Tree of Life. eLife. 2023;12:RP88251.

Miller MA, Pfeiffer W, Schwartz T. Creating the CIPRES Science Gateway for inference of large phylogenetic trees. In: 2010 gateway computing environments workshop (GCE). Ieee, pp. 1-8; 2010.

Moya-Laraño J, Vinković D, De Mas E, Corcobado G, Moreno E. Morphological evolution of spiders predicted by pendulum mechanics. PLoS ONE. 2008;3:e1841.

Noss RF. Indicators for monitoring biodiversity: a hierarchical approach. Conserv Biol. 1990;4:355–364.

Paradis E, Schliep K. ape 5.0: an environment for modern phylogenetics and evolutionary analyses in R. Bioinformatics. 2019;35:526–528.

Pavoine S, Bonsall MB. Measuring biodiversity to explain community assembly: a unified approach. Biol Rev. 2011;86:792–812.

Pereira HM, Belnap J, Brummitt N, Collen B, Ding H, Gonzalez-Espinosa M, et al. Global biodiversity monitoring. Front Ecol Environ. 2010;8:459–460.

Pereira HM, Ferrier S, Walters M, Geller GN, Jongman RHG, Scholes RJ, et al. Essential biodiversity variables. Science. 2013;339:277–278.

Petchey OL, Gaston KJ. Functional diversity (FD), species richness and community composition. Ecol Lett. 2002;5:402–411.

Ricotta C, Bacaro G, Marignani M, Godefroid S, Mazzoleni S. Computing diversity from dated phylogenies and taxonomic hierarchies: does it make a difference to the conclusions? Oecologia. 2012;170:501–506.

Rodrigues AS, Gray CL, Crowter BJ, Ewers RM, Stuart SN, Whitten T, Manica A. A global assessment of amphibian taxonomic effort and expertise. BioScience. 2010;60(10):798–806.

Schilthuizen M, Vairappan CS, Slade EM, Mann DJ, Miller JA. Specimens as primary data: museums and ‘open science’. Trends Ecol Evol. 2015;30:237–238.

Smith SA, O’Meara BC. treePL: divergence time estimation using penalized likelihood for large phylogenies. Bioinformatics. 2012;28:2689–2690.

Stamatakis A. RAxML version 8: a tool for phylogenetic analysis and post-analysis of large phylogenies. Bioinformatics. 2014;30:1312–1313.

Stork NE, Samways MJ, Eeley HAC. Inventorying and monitoring biodiversity. Trends Ecol Evol. 1996;11:39–40.

Tuomisto H. A diversity of beta diversities: Straightening up a concept gone awry. Part 1. Defining beta diversity as a function of alpha and gamma diversity. Ecography. 2010;33:2–22.

Upham NS, Esselstyn JA, Jetz W. Inferring the mammal tree: species-level sets of phylogenies for questions in ecology, evolution, and conservation. PLoS Biol. 2019;17:e3000494.

Van Klink R, Bowler DE, Gongalsky KB, Swengel AB, Gentile A, Chase JM. Meta-analysis reveals declines in terrestrial but increases in freshwater insect abundances. Science. 2020;368:417–420.

Van Klink R, August T, Bas Y, Bodesheim P, Bonn A, Fossøy F, et al. Emerging technologies revolutionise insect ecology and monitoring. Trends Ecol Evol. 2022;37:872– 885.

Van Swaay C, Regan E, Ling M, Bozhinovska E, Fernandez M, Marini-Filho OJ, et al. Guidelines for Standardised Global Butterfly Monitoring. Group on Earth Observations Biodiversity Observation Network, Leipzig, Germany. GEO BON Technical Series 1, 32pp; 2015.

Wauchope HS, Amano T, Sutherland WJ, Johnston A. When can we trust population trends? A method for quantifying the effects of sampling interval and duration. Methods Ecol Evol. 2019;10:2067–2078.

Wheeler WC, Coddington JA, Crowley LM, Dimitrov D, Goloboff PA, Griswold CE, et al. The spider tree of life: phylogeny of Araneae based on target-gene analyses from an extensive taxon sampling. Cladistics. 2016;33:574–616.

White ER. Minimum time required to detect population trends: The need for long-term monitoring programs. BioScience. 2019;69:40–46.

Wilman H, Belmaker J, Simpson J, de la Rosa C, Rivadeneira MM, Jetz W. EltonTraits 1.0: SpeciesLJlevel foraging attributes of the world’s birds and mammals: Ecological Archives E095LJ178. Ecology. 2014;95(7):2027–2027.

World Spider Catalog (2024). World Spider Catalog. Version 24.5. Natural History Museum Bern, online at http://wsc.nmbe.ch, accessed on 14.01.2024. 10.24436/2

Yoccoz NG, Nichols JD, Boulinier T. Monitoring of biological diversity in space and time. Trends Ecol Evol. 2001;16:446–453.

