## Appendix S1 for "Optimal inventorying and monitoring of taxonomic, phylogenetic and functional diversity"

**Table S1. Eighteen sampled sites in Iberian oak forests.**

| **Site** | **Latitude** | **Longitude** | **Elevation (m)** | **Reference** |
| --- | --- | --- | --- | --- |
| **Aiguestortes 1** | 42.5496 | 0.8725 | 1760 | Crespo et al. 2018 |
| **Aiguestortes 2** | 42.5491 | 0.8714 | 1740 | Crespo et al. 2018 |
| *** Arrábida** | 38.5067 | -8.9752 | 60 | Cardoso et al. 2008a |
| **Cabañeros 1** | 39.3566 | -4.3591 | 760 | Crespo et al. 2018 |
| **Cabañeros 2** | 39.3516 | -4.3589 | 740 | Crespo et al. 2018 |
| **Cabañeros 3** | 39.3618 | -4.4173 | 770 | Crespo et al. 2018 |
| **Cabañeros 4** | 39.3634 | -4.4170 | 770 | Crespo et al. 2018 |
| *** Gerês** | 41.7953 | -8.1363 | 660 | Cardoso et al. 2008b |
| **Monfragüe 1** | 39.8330 | -6.0641 | 320 | Crespo et al. 2018 |
| **Monfragüe 2** | 39.8280 | -6.0325 | 320 | Crespo et al. 2018 |
| **Ordesa 1** | 42.6068 | 0.1313 | 1400 | Crespo et al. 2018 |
| **Ordesa 2** | 42.5943 | 0.1529 | 1160 | Crespo et al. 2018 |
| **Picos de Europa 1** | 43.1445 | -4.9267 | 1070 | Crespo et al. 2018 |
| **Picos de Europa 2** | 43.1777 | -4.9058 | 760 | Crespo et al. 2018 |
| **Picos de Europa 3** | 43.1435 | -4.9488 | 1100 | Crespo et al. 2018 |
| **Picos de Europa 4** | 43.1723 | -4.9086 | 940 | Crespo et al. 2018 |
| **Sierra Nevada 1** | 36.9615 | -3.4188 | 1790 | Crespo et al. 2018 |
| **Sierra Nevada 2** | 37.1838 | -3.2628 | 1710 | Crespo et al. 2018 |

At two sites (with *) 320 samples were performed and these were used only for the optimization of inventorying. At 16 sites, 24 samples were performed, and these were used only for the optimization of monitoring.
